# Linking anterior cingulate sulcal variability to attention-deficit hyperactivity disorder behavior

**DOI:** 10.64898/2026.10.12.758584

**Authors:** Robin F. Hadveck, Abigail Paras, Megan Anctil, Enrique Martinez, Charley M. Gallardo, Samira A. Maboudian, Ethan H. Willbrand, Kevin S. Weiner

**Affiliations:** Department of Psychology, University of California, Berkeley, Berkeley, CA, USA; Helen Wills Neuroscience Institute, University of California, Berkeley, Berkeley, CA, USA; Department of Neuroscience, University of California, Berkeley, Berkeley, CA, USA; Medical Scientist Training Program, University of Wisconsin School of Medicine and Public Health, Madison, USA; Department of Radiology, University of Wisconsin School of Medicine and Public Health, Madison, USA; Neuroscience Training Program, University of Wisconsin School of Medicine and Public Health, Madison, USA

**Author notes:** Corresponding Author Kevin S. Weiner. co-first authors (Robin F. Hadveck, Abigail Paras).

**Keywords:** ADHD, Cingulate cortex, Impulsivity, Neuroanatomy, Neuroimaging, Psychopathology

## Abstract

**Background:** The anterior cingulate cortex (ACC) is consistently implicated in attention-deficit/hyperactivity disorder (ADHD). ACC variability, including its most variable structure, the paracingulate sulcus (PCGS), has been known since the late 1800s, and since then, has been linked to cognitive control, impulsivity, and psychiatric conditions comorbid with ADHD, including autism spectrum disorder and schizophrenia. Yet, the role of the PCGS in ADHD remains unexamined at the individual level.

**Methods:** We characterized PCGS incidence, asymmetry, and morphology in 247 children and adolescents (144 ADHD: 10.9 ± 2.73 years old, 75% male; 103 neurotypical: 12.2 ± 3.17 years old, 50% male) from the ADHD-200 dataset. The PCGS was manually identified in FreeSurfer cortical surface reconstructions. Morphometric measures, including sulcal length, depth, and cortical thickness, were extracted and normalized to hemisphere-wide variables. Between-group analyses tested whether PCGS presence, asymmetry, and quantitative morphology differed by diagnosis, while behavioral analyses examined associations with inattention, hyperactivity-impulsivity, and cognitive measures. Age and gender were included as covariates.

**Results:** Analyses revealed that, in both NT and ADHD individuals, there was significant leftward asymmetry in PCGS presence. Between groups, the PCGS was less likely to be present in individuals with ADHD in the left hemisphere. Leftward PCGS absence was also associated with significantly higher ADHD index and hyperactivity-impulsivity scores.

**Conclusions:** These findings advance understanding of how early neurodevelopmental variation in cortical folding may serve as a structural intermediary between genetic liability and the behavioral expression of ADHD.

## Introduction

Identifying links between individual differences in brain structure and function and how that relates to individual differences in cognition across the lifespan and disorder is a major goal in basic and translational neuroscience. In terms of brain structure, there are many features to measure, including but not limited to, microarchitecture, cortical folding, connectivity, subcortical and cerebellar volumes, and an oft-overlooked feature: tertiary sulcal morphology. Tertiary sulci appear last in gestation, continue to develop after birth, and have been linked to cognition across the lifespan and in different clinical populations (1–12). Notably, some tertiary sulci can be identified in every hemisphere, while others are variable across species and even between hemispheres in the same person (2,5,13–20).

For example, in the anterior cingulate cortex (ACC), the paracingulate sulcus (PCGS) is highly variable from hemisphere to hemisphere. The PCGS, which is a hominoid-specific structure, is present in at least one hemisphere 60-70% of the time with a leftward asymmetry (1,2,14,21–25). The presence or absence of the PCGS affects the underlying microarchitecture of the ACC (21,26) and often exhibits greater leftward asymmetry in males than females (22,23,27–29). Importantly, PCGS presence/absence and asymmetry are cognitively and clinically consequential: its presence and lateralization have been linked to executive function, inhibitory control, and trait impulsivity, as well as altered in different disorders such as schizophrenia, obsessive-compulsive disorder, autism spectrum disorder, and bipolar disorder (2,30–38). However, an unaddressed question is: Is the hemispheric presence/absence or asymmetry of the PCGS in individuals with attention-deficit/hyperactivity disorder (ADHD) related to individual differences in cognition and behavior?

To fill these gaps in knowledge, the present study sought to (i) characterize PCGS incidence and morphology in ADHD individuals, (ii) assess whether PCGS incidence and morphology differs in ADHD individuals compared to NT individuals, and (iii) test if PCGS presence/absence is related to ADHD-related behaviors. Given previous findings in other disorders involving folding of the ACC, we hypothesize that hemispheric PCGS presence and left-right asymmetry differs in ADHD and that paracingulate presence and asymmetry will be related to the ADHD-related behaviors of inattention, hyperactivity, and/or impulsivity. Importantly, findings from this approach could inform a converging body of research proposing a hemispheric asymmetry hypothesis to characterize neurodevelopmental differences in ADHD (39–44). This hypothesis specifically suggests that atypical development of the brain networks that underlie core ADHD symptomatology appear as alterations in the asymmetry of structural and functional neuroimaging measures. Thus, the variable presence and asymmetry of the PCGS makes it an ideal target to further investigate this hypothesis.

Here, we characterized PCGS incidence, asymmetry, and morphology in 247 children and adolescents separately in each hemisphere (144 ADHD: 10.9 ± 2.73 years old, 75% male; 103 neurotypical: 12.2 ± 3.17 years old, 50% male) drawn from the ADHD-200 dataset. In both NT and ADHD individuals, there was a significant leftward asymmetry in PCGS presence, an asymmetry that is consistent with previous findings. The PCGS was less likely to be present in individuals with ADHD, specifically in the left hemisphere. Further, the presence of the PCGS in the left hemisphere was associated with lower ADHD index, as well as inattentive and hyperactivity-impulsivity scores. These findings advance understanding of how early neurodevelopmental variation in cortical folding may serve as a structural intermediary between genetic liability and the behavioral expression of ADHD. They also directly inform the recent hemispheric asymmetry hypothesis by demonstrating that a specific, variable sulcus in ACC contributes to individual differences in ADHD symptomatology.

## Methods and Materials

### Participants

Participant data was obtained from the publicly available ADHD-200 dataset (https://fcon_1000.projects.nitrc.org/indi/adhd200/). Study procedures were approved by each site’s local Institutional Review Board and parental written informed consent and child assent were obtained from all participants. We specifically leveraged data from 247 participants from the New York University Child Study Center site, which consisted of 103 NT participants (7.17–18.0 years old; 50.49% female) and 144 participants with ADHD (7.24–17.6 years old; 24.48% female). Psychiatric diagnoses were based on evaluations with the Schedule of Affective Disorders and Schizophrenia for Children-Present and Lifetime Version (KSADS-PL) administered to parents and children and the Conners’ Parent Rating Scale-Revised, Long version (CPRS-LV). Intelligence was evaluated with the Wechsler Abbreviated Scale of Intelligence (WASI). Inclusion in the ADHD group required a diagnosis of ADHD based on parent and child responses to the KSADS-PL as well as on a T-score greater than or equal to 65 on at least one ADHD related index of the CPRS-LV. Psychostimulant drugs were withheld at least 24 hours before scanning. Inclusion criteria for NT participants required absence of any Axis-I psychiatric diagnoses per parent and child KSADS-PL interview, as well as T-scores below 60 for all the CPRS-LV ADHD summary scales. Estimates of Full Scale Intelligence Quotient (FSIQ) above 80, right-handedness and absence of other chronic medical conditions were required for all participants. More detailed inclusion/exclusion criteria and demographic information of the participants can be found at the ADHD-200 website. Demographic, diagnostic, and behavioral details of this sample are provided in **Table 1**.

**Table 1.**
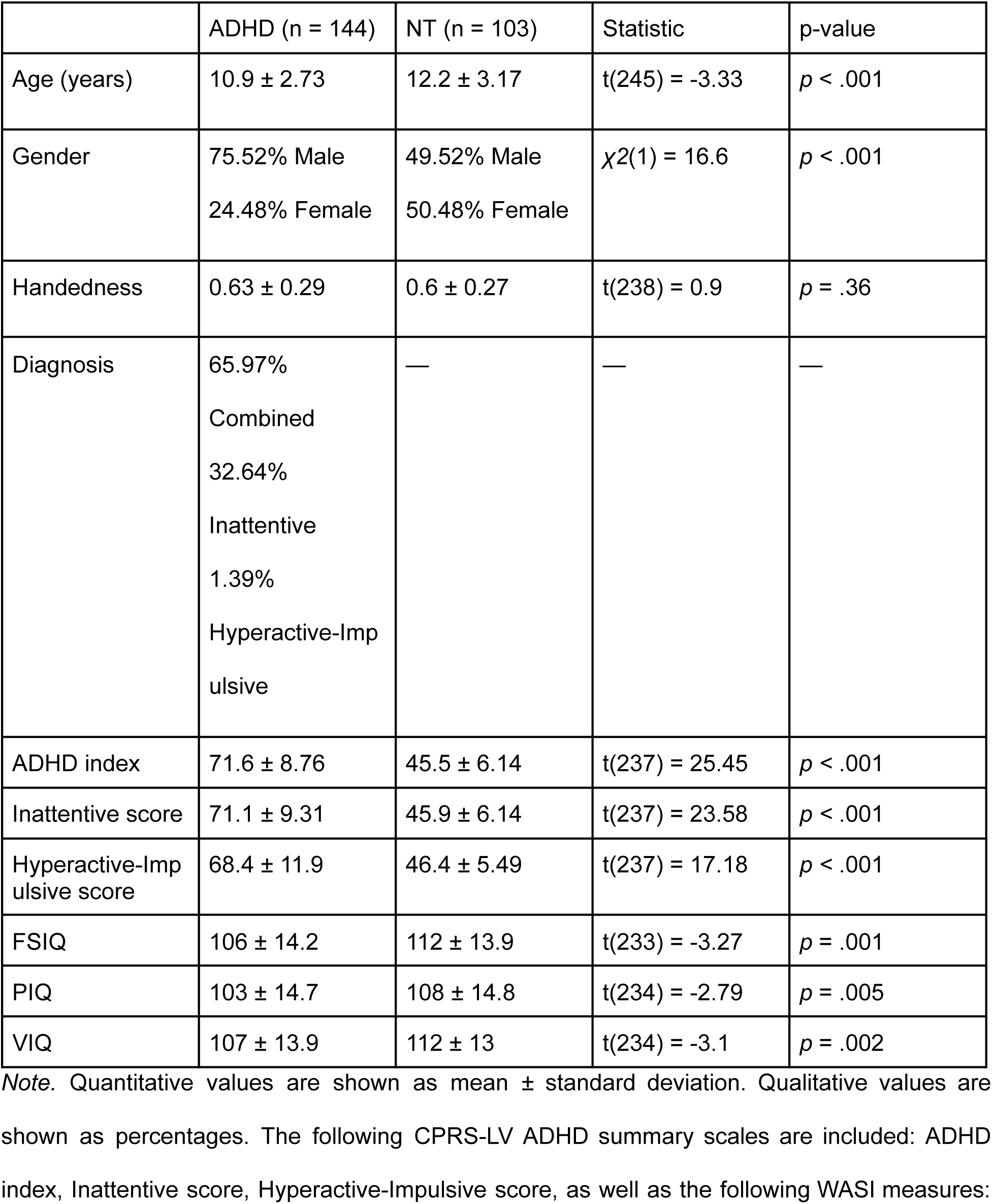
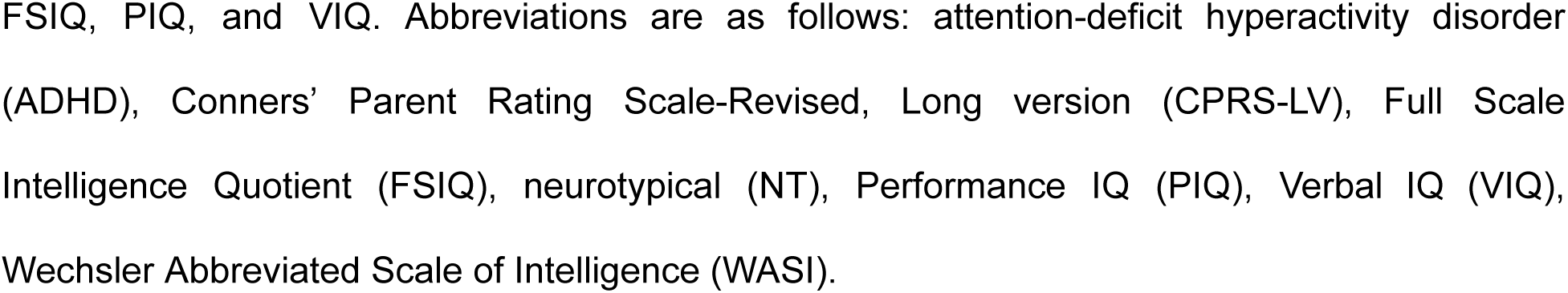
Demographic and behavioral data for each diagnostic group.

|  | ADHD (n = 144) | NT (n = 103) | Statistic | p-value |
| --- | --- | --- | --- | --- |
| Age (years) | 10.9 ± 2.73 | 12.2 ± 3.17 | t(245) = -3.33 | p < .001 |
| Gender | 75.52% Male<br>24.48% Female | 49.52% Male<br>50.48% Female | $\chi^2(1) = 16.6$ | p < .001 |
| Handedness | 0.63 ± 0.29 | 0.6 ± 0.27 | t(238) = 0.9 | p = .36 |
| Diagnosis | 65.97%<br>Combined<br>32.64%<br>Inattentive<br>1.39%<br>Hyperactive-Impulsive | — | — | — |
| ADHD index | 71.6 ± 8.76 | 45.5 ± 6.14 | t(237) = 25.45 | p < .001 |
| Inattentive score | 71.1 ± 9.31 | 45.9 ± 6.14 | t(237) = 23.58 | p < .001 |
| Hyperactive-Impulsive score | 68.4 ± 11.9 | 46.4 ± 5.49 | t(237) = 17.18 | p < .001 |
| FSIQ | 106 ± 14.2 | 112 ± 13.9 | t(233) = -3.27 | p = .001 |
| PIQ | 103 ± 14.7 | 108 ± 14.8 | t(234) = -2.79 | p = .005 |
| VIQ | 107 ± 13.9 | 112 ± 13 | t(234) = -3.1 | p = .002 |

*Note.* Quantitative values are shown as mean ± standard deviation. Qualitative values are shown as percentages. The following CPRS-LV ADHD summary scales are included: ADHD index, Inattentive score, Hyperactive-Impulsive score, as well as the following WASI measures:
FSIQ, PIQ, and VIQ. Abbreviations are as follows: attention-deficit hyperactivity disorder (ADHD), Conners' Parent Rating Scale-Revised, Long version (CPRS-LV), Full Scale Intelligence Quotient (FSIQ), neurotypical (NT), Performance IQ (PIQ), Verbal IQ (VIQ), Wechsler Abbreviated Scale of Intelligence (WASI).

### Neuroanatomical data

#### Imaging Acquisition

Standard T1-weighted MPRAGE anatomical scans (TR = 2530 ms, TE = 3.25 ms, 1.3 × 1 × 1.3 mm^3^ voxels) were acquired for cortical morphometric analyses in these participants using a 3T SIEMENS MAGNETOM Allegra syngo MR scanner at New York University Langone Medical Center.

#### Cortical Surface Reconstruction

Cortical surface reconstructions were generated for each participant from their T1-weighted image using the standard FreeSurfer pipeline (v6.0) (45–47). The manual identification of the PCGS was performed on each cortical surface in native space.

#### Paracingulate Sulcus Identification, Classification, and Morphometry

To characterize anterior cingulate sulcal organization, we evaluated each hemisphere for the presence of the PCGS, a tertiary fold located dorsal to and running roughly parallel with the CGS (22,23). This assessment followed commonly used criteria in the PCGS literature (1,24,37,38,48,49). Specifically, the PCGS was coded as “present” when a horizontally oriented segment extended alongside the CGS for ≥20 mm and reached a minimum depth of 4 mm. Hemispheres lacking any segment meeting these thresholds were labeled “absent.” This binary approach is consistent with prior work and avoids sub-categories such as “prominent,” which show inconsistent reliability and demographic sensitivity (2,23,27–29,48,50). We also described left–right asymmetries using established terminology (22,23). A hemisphere pair was considered asymmetric when only one side met the criteria for PCGS presence; conversely, hemispheres were labeled symmetric when the sulcus was either present or absent on both sides. PCGS coding was performed by trained raters, blind to group. To evaluate interrater reliability, 15% of hemispheres were labeled independently, which yielded near-perfect agreement (Cohen’s κ = 0.90). Representative examples of presence, absence, and asymmetry are shown in **Figure 1**.

**Figure 1.**
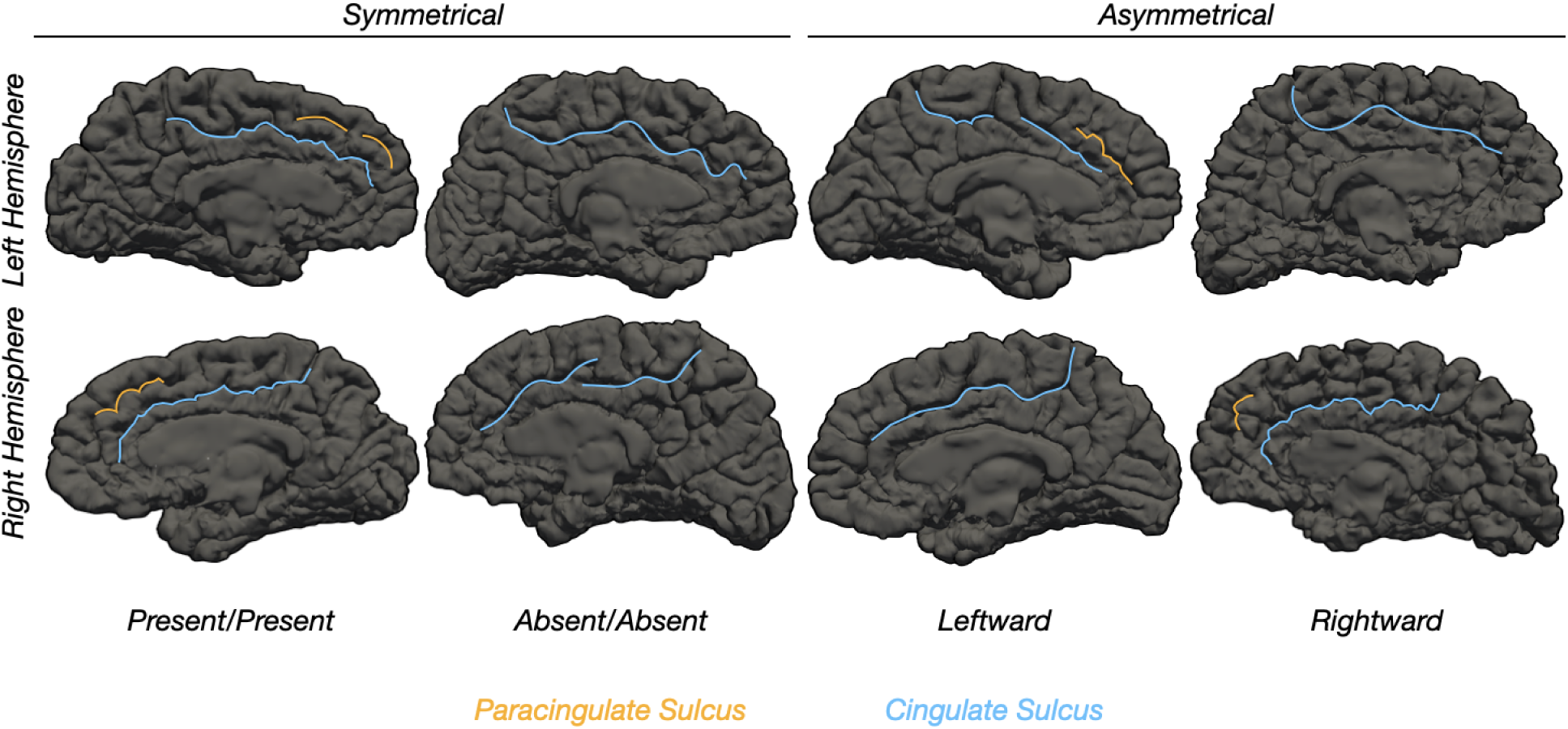
Examples of PCGS sulcal patterning. Sulcal patterning is defined by hemispheric presence/absence of the PCGS, which runs posterior and parallel to the CGS when present (PCGS: orange; CGS: blue). The images depict the inter-hemispheric asymmetry/symmetry (columns) and intra-hemispheric presence/absence (rows) of the PCGS in four example participants (columns) on the pial FreeSurfer cortical surface reconstructions. Sulci are black, and gyri are dark gray. Abbreviations are as follows: cingulate sulcus (CGS), paracingulate sulcus (PCGS).

For morphometric quantification, the PCGS was manually delineated on each hemisphere as a FreeSurfer .label file using tksurfer. Sulcal length was computed as the maximum geodesic distance between boundary vertices using methods implemented in the pycortex package (51), with discontinuous sulcal segments merged and gyral interruptions excluded. Depth measurements were derived from the fundus to the pial surface using a modified procedure adapted from the standard FreeSurfer workflow (52). Another standard morphological metric, specifically, mean cortical thickness (CT), was extracted with mris_anatomical_stats (53). All quantitative morphological variables were normalized to the relevant hemisphere-wide variable (hemispheric surface area, hemispheric max depth, and hemispheric CT mean).

### Quantification and Statistical Analysis

All statistical tests were implemented in R (v4.1.2). All P-values were adjusted using the Benjamini-Hochberg false discovery rate (FDR) procedure. Statistical significance was defined as FDR-adjusted P < 0.05. We conducted three series of analyses as follows.

#### Within-group analyses

For each diagnostic group separately, we tested for intra- and inter-hemispheric differences in PCGS presence and PCGS asymmetry. Differences in categorical variables were tested using chi-squared tests. McNemar’s test for symmetry was used to specifically assess PCGS asymmetry. We assessed whether gender, age, and handedness related to left/right PCGS presence and asymmetry with binomial logistic regression and general linear models (GLMs) for PCGS patterning [0 (present or symmetric), 1 (absent or asymmetric)] with gender, age, and handedness as the primary factors. GLM effect size was quantified as the Odds ratio (OR).

#### Between-group analyses

We next tested whether PCGS features were related to diagnosis. We first ran binomial logistic regression and general linear models (GLMs) for diagnostic group [0 (NT), 1 (ADHD)] with left/right PCGS presence and PCGS asymmetry as the primary factors, respectively. We then ran a GLM with the primary factors instead being the four quantitative morphological features of interest (left/right PCGS length, left/right PCGS depth, and left/right PCGS CT mean). GLM effect size was quantified as the OR. In all analyses, age, gender, and handedness were included as control variables given the group differences (**Table 1**).

#### Behavioral analyses

We finally tested whether ADHD-related behaviors were associated with the aforementioned PCGS features using the CPRS-LV scores (**Table 1**). For each CPRS-LV test, associations with qualitative PCGS features were tested using an analysis of variance (ANOVA) and associations with quantitative PCGS features were compared using linear regressions. Effect sizes were quantified as partial eta-squared (η_p_ ^2^) and adjusted R^2^, respectively. We included age, gender, and handedness as control variables. In the quantitative morphological analyses, the models all controlled for the relevant hemisphere-wide variable (hemispheric surface area, max depth, and CT mean).

#### Meta-analysis

To test potential relationships between the PCGS and the broad literature on dmPFC and dACC functions related to ADHD, we downloaded the uniformity test maps for the following three terms from NeuroSynth (55): (i) *conflict* (337 studies), (ii) *value* (470 studies), and (iii) *error* (464 studies). Each uniformity test map was generated from a *χ2* test (FDR-corrected, P = 0.01) comparing the activation in each voxel for studies containing the term compared with what one would expect if activation were uniformly distributed throughout the gray matter. Each map was projected to *fsaverage* surface space with the *mri_vol2surf* FreeSurfer function to spatially relate them to PCGS probabilistic maps in each hemisphere, from Willbrand *et al*. (38).

## Results

### Significant variability in presence of PCGS within and between hemispheres

Consistent with previous studies in NT and clinical neurodevelopmental samples (2), we observed significant variability in presence of PCGS within and between hemispheres (**Fig. 2**). In both groups, the PCGS was more often present than absent in the left hemisphere (NT: χ2(1) = 36.13, P = 0.000000001; ADHD: χ2(1) = 13.44, P = 0.0002) but not the right hemisphere (NT: χ2(1) = 1.64, P = 0.20; ADHD: χ2(1) = 0.44, P = 0.50; **Fig. 2**). The presence of a PCGS was more frequent in the left than right hemisphere in NT participants (χ2(1) = 12.84, P = 0.0003), and in ADHD participants as well (χ2(1) = 4.65, P = 0.031; **Fig. 2**). In addition, implementing McNemar’s test for symmetry revealed significant asymmetry in PCGS presence in both NT participants and ADHD participants (NT: χ2(1) = 11.08, P = 0.0008; ADHD: χ2(1) = 6.00, P = 0.014; **Fig. 2**).

**Figure 2.**
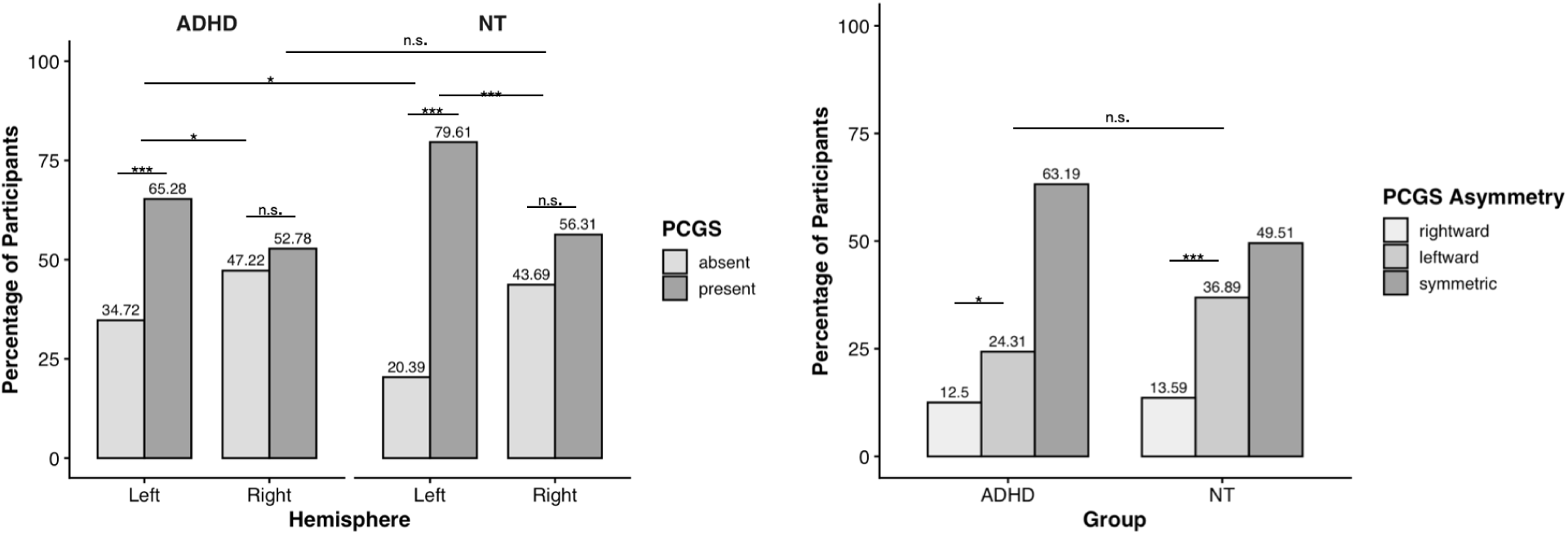
Individuals with ADHD are less likely to have a PCGS in the left hemisphere. Left: Bar plot visualizing PCGS incidence as a function of hemisphere (x-axis), group (NT=103; ADHD=144; Methods), and presence (colors; see key). Right: Bar plot visualizing PCGS incidence as a function of group (x-axis) and asymmetry (colors; see key). The upper asterisk denotes GLM results for between-group comparisons of asymmetry, while lower asterisks depict McNemar test results for within-group comparisons of asymmetry. Percentages are included above each column. Abbreviations are as follows: Attention Deficit Hyperactivity Disorder (ADHD), group, neurotypical (NT), paracingulate sulcus (PCGS). Asterisks indicate the following p-value thresholds: *** p < 0.001, * p < 0.05, n.s. p > .05.

### Individuals with ADHD are less likely to have a PCGS in the left hemisphere

Next, we explicitly compared PCGS presence and asymmetry between the two groups by implementing logistic regression GLMs with a predictor of group (controlling for age, gender, and handedness). There was a main effect of group on left PCGS presence, indicating that ADHD participants were less likely than NT participants to have a PCGS in the left hemisphere (Left: P = 0.021; Right: P = 0.67; **Fig. 2**). Moreover, there was a marginal main effect of group on PCGS asymmetry (P = 0.073), which showed that NT participants had lower odds of having a symmetric PCGS compared to ADHD participants (OR = 0.49 [0.25-0.93]; **Fig. 2**). We then examined whether quantitative morphological features of the PCGS (length, depth, and CT mean and CT STD) differed between the two groups by implementing LMEs with predictors of group and hemisphere (controlling for age, scanner site, and the respective hemispheric-level morphological variable). There were no significant main effects of group or group x hemisphere interactions for any of the metrics considered (all Ps > 0.05). There were no other hemispheric differences observed for the other three quantitative morphological features (all Ps > 0.05).

### Hemispheric PCGS Presence Correlates with Behavioral Scores

Building on our qualitative and quantitative neuroanatomical analyses, we then related PCGS presence in each hemisphere separately to each of three behavioral metrics: ADHD index, inattentive score, and hyperimpulsivity. A Type III analysis of variance (ANOVA) with factors of LH and RH PCGS presence revealed a main effect of left PCGS presence for two behavioral metrics beyond what is explained by other factors such as age, gender, and handedness, in which LH PCGS absence was related to an increased severity of each score or index (ADHD index: F1,226, P = 0.049, ηp2 = 0.017; Hyperimpulsivity score: F1,226, P = 0.021, ηp2 = 0.023). Impulsivity was marginal (F1,226, P = 0.073, ηp2 = 0.014; **Fig. 3**).

**Figure 3.**
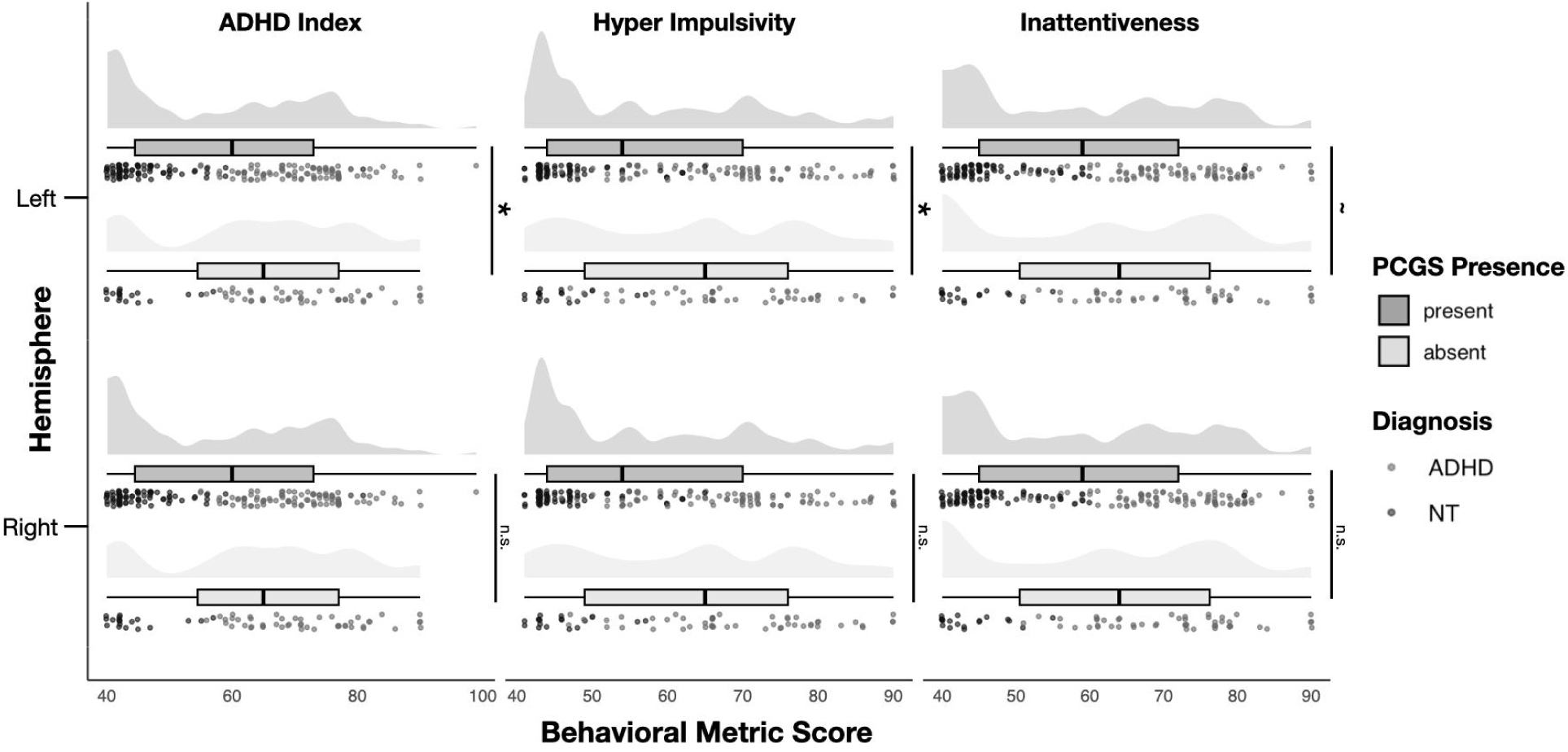
Leftward PCGS absence is associated with higher ADHD index and hyperactivity-impulsivity scores. Raincloud plots showing ADHD index (left), hyperactivity-impulse scores (middle), and inattentive scores (right) as a function of PCGS presence/absence in each hemisphere. Data are shown for the total sample (N=247, ADHD n=144, NT n=103). Half-violin density clouds depict sample distributions. Boxplots indicate mean and interquartile ranges, with individual data points (‘rain’) jittered by the diagnostic group (light gray: ADHD; dark gray: NTs). Significance bars depict Type III ANOVA main effects without diagnostic interaction. *p < .05; ∼p<.08, n.s. p > .08

Group differences for all three behavioral metrics were also significantly driven by age (ADHD index: F1,226, P = 0.001, ηp2 = 0.044; Inattentive scores: F1,266, P < 0.001, ηp2 = 0.047; Hyperimpulsivity score: F1,226, P < 0.001, ηp2 = 0.014) and in the case of Hyperimpulsivity, gender (F1,226, P = 0.042, ηp2 = 0.018). To test if this brain-behavior relationship was modified by clinical diagnostic status, we introduced the interaction term of diagnosis. None of the interactive terms demonstrated significance (Ps > 0.05). Additionally, while nominal relationships were observed, no quantitative PCGS features had a significant relationship with any behavioral metric (**Supplementary Materials**). When further assessing the relationship among quantitative morphological features (PCGS length, depth, and cortical thickness) relative to these indices and scores, no comparisons survived when adjusting for False Discovery Rate (**Supplementary Materials**).

## Discussion

The present study sought to examine (i) PCGS incidence and morphology in ADHD individuals, as well as comparisons relative to NTs, and (ii) whether the presence or absence of the PCGS contributed to ADHD-related behaviors. The results of this approach identify a specific, lateralized, and behaviorally relevant PCGS variability to individual differences in cognition, behavior, and psychopathology. These findings support the proposed hemispheric asymmetry hypothesis to characterize neurodevelopmental differences in ADHD (39–44). In the sections below, we discuss these findings in the context of (i) the role of the PCGS in the function of dorsal medial prefrontal cortex (dmPFC) and the dorsal anterior cingulate cortex (dACC), (ii) underlying neuroanatomical mechanisms related to the presence or absence of the PCGS, and (iii) behavioral implications of the development and maturation of the PCGS in ADHD.

### The role of the PCGS in the function of dorsal medial prefrontal cortex (dmPFC) and the dorsal anterior cingulate cortex (dACC)

A recent synopsis of functional parcellations of dmPFC and dACC proposes that a contributing reason for an "ill-defined brain region" in this cortical expanse is the presence or absence of the PCGS. In particular, Clairis and Lopez-Persem synthesize previous work and consider different functions in this concentrated cortical area such as value, conflict monitoring, and error-likelihood while considering different clinical implications, including ADHD (54). Here, in order to further examine this relationship at the meta-analytic level, we examined the relationship between the maximum probability of the PCGS and relevant search terms in NeuroSynth (55). When indexing the term *conflict* in Neurosynth, the activated clusters across 337 studies overlap with the maximum probability map (MPM) of the PCGS (**Fig. 4A**).

**Figure 4.**
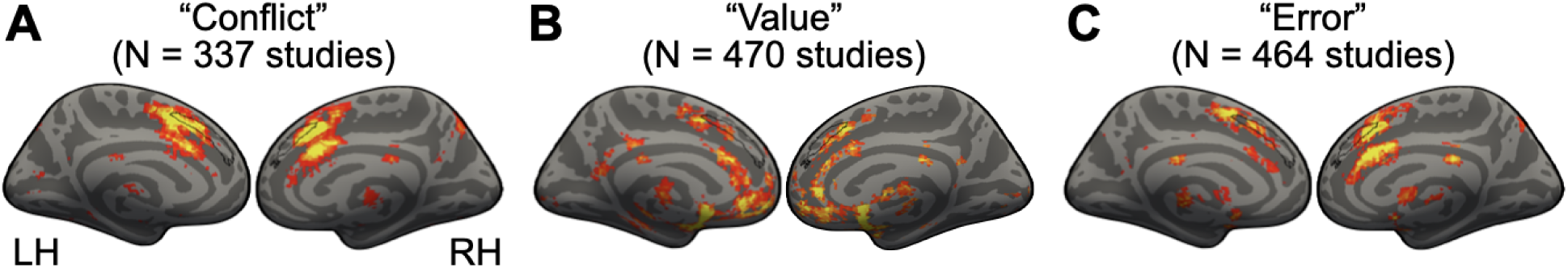
The PCGS co-localizes with Neurosynth meta-analytic clusters related to conflict, value, and error terms. **A.** Left and right hemisphere inflated *fsaverage* cortical surfaces (left surface: left hemisphere, LH; right surface: right hemisphere, RH) displaying overlap of a whole-brain FDR-corrected (P = 0.01) uniformity-test meta-analysis z-score map of the “conflict” term (downloaded from Neurosynth; https://neurosynth.org/) (55) with a maximum probability map (25% participant overlap threshold) of the PCGS (black outline) (38). **B.** Same as **A.**, but for the “value” term. **C.** Same as **A.**, but for the “error” term.

Additionally, the activated clusters across 470 studies for *value* (**Fig. 4B**) and 464 studies for *error* (**Fig. 4C**) overlap with the MPM of the PCGS. While this meta-analytic approach complements and extends the present findings, as well as aligns with the proposal by Clairis and Lopez-Persem (54), it also motivates future studies to examine the relationship among functional activations (during conflict monitoring, value, or error-likelihood tasks), connectivity (both anatomical and functional), and the presence/absence of the PCGS in individual participants with a precision imaging approach.

### Including the presence or absence of sulci in future ADHD studies: From mechanisms to subtypes

While (to our knowledge) this is the first study to examine how the presence or absence of the PCGS is related to ADHD symptoms in individual participants, prior studies implementing meta-analyses and large-scale studies revealed differences between brain morphology in ADHD individuals compared to age-matched controls. For example, individuals with ADHD exhibit differences in sulcal pit (e.g., deep sulcal points, which we discuss further in the next section) depths, cortical thickness, gray matter and white matter volume, and cortical surface area, particularly within prefrontal and anterior cingulate cortices (56–60). Furthermore, sulcal depth in the parietal, temporal, and occipital regions, in addition to increased gyrification of frontal cortical regions, have been shown to predict increased treatment response to ADHD medication (61,62). As there are additional variable tertiary sulci (aside from the PCGS) in these association cortices implicated in previous studies, investigating if the presence of sulci in these cortical expanses of ADHD individuals could provide further clinical insights into the neuroanatomical correlates of ADHD.

In terms of mechanism, the presence or absence of the PCGS influences the underlying microarchitecture, and in turn, local cortical microcircuits, as well as network properties of long-range cortical networks. Indeed, previous findings show that PCGS absence indicates alterations in the functional and cytoarchitectonic organization of ACC, and as a result, different connectivity patterns and microarchitecture contributing to differences in function (21,26,63). Recent findings further support this PCGS-connectivity relationship. Danstrom and colleagues showed that white matter connectivity underlying the ACC may be altered in the presence of a PCGS, particularly with respect to bilateral connectivity, which is modulated by left-hemisphere PCGS morphology (64). Further, this structural influence is consistent with findings in which disrupted ACC white matter integrity is accompanied by increased fluctuations in spontaneous brain activity in ADHD (65).

An interesting avenue for future research would be to test if the presence/absence of the PCGS is more strongly related to different subtypes of ADHD (66–68), and also, if the presence/absence of the PCGS predicts differences of white matter connectivity and microarchitecture among these different subtypes. Recent findings show promise for this future endeavor: ADHD subtypes display distinct connectivity profiles in which the combined subtype shows higher nodal degree in the anterior cingulate than controls, without significant gray matter volume or network organizational differences (69). Findings from these future studies could further inform the hemispheric asymmetry hypothesis of ADHD using multimodal data, from connectivity to microarchitecture in-vivo (for example, using quantitative MRI).

### PCGS: Maturation and behavioral implications in ADHD

Future studies can also further explore how developmental and morphological alterations of dmPFC and dACC are impacted by the presence or absence of the PCGS and in turn, how that influences ADHD severity or subtypes. For instance, dmPFC maturation toward the cingulate gyrus is delayed in ADHD by a median of 1.9 years (70,71). In terms of sulcal morphology, sulcal pits that reflect cortical folding maturation are reduced in ADHD compared to NTs, particularly across the bilateral cingulate sulcus (59). These sulcal pit differences directly impact the dynamics of paracingulate gyrus folding, are under strong genetic control, and remain tightly coupled with functional cortical organization (59). Nevertheless, prior research on the PCGS shows weak additive genetic factors influencing the presence of a PCGS in siblings, but a significant relationship of twin-status relating PCGS organization and individual differences in behavior (spanning motor, affective, and cognitive tasks) (1).

In terms of behavior, the present study expanded on PCGS variability in cognitive control, impulsivity, and psychiatric and neurodevelopmental conditions. Left PCGS presence was associated with greater symptom severity, which is a finding that builds on previous work that individuals with leftward PCGS patterns have heightened impulsivity, a related transdiagnostic trait (38). The present results also extend earlier findings demonstrating reduced PCGS asymmetry in individuals with autism spectrum disorder (ASD) relative to NT individuals (37), a condition with significant diagnostic overlap with ADHD (72). Considered in conjunction with that prior finding, the present results suggest that while both ADHD and ASD neurodevelopment are associated with atypical PCGS patterning relative to NT individuals, there is a distinct difference: ASD is characterized by disrupted hemispheric asymmetry while ADHD has reduced categorical presence of the left PCGS with no change in overall asymmetry. This subtle distinction nurtures the possibility that PCGS related phenotypes may act as a marker for partially dissociable neurodevelopmental pathways across comorbid conditions, a possible direction for future studies with transdiagnostic samples. Additionally, future studies modeling ADHD both as a categorical diagnosis and as a continuous, dimensional spectrum of symptoms, would help determine whether PCGS-behavior associations are specific to clinical diagnosis or extend across the full range of attention and impulse-control variation in the general population. Given that ADHD symptoms and their neural correlates change across development, longitudinal designs (as opposed to the cross-sectional design implemented here) tracking the same individuals would help determine whether the relationship between leftward asymmetry and symptom severity is stable, or whether it strengthens or attenuates with age as prefrontal-cingulate circuits mature.

On a broader scale, these findings are consistent with altered PCGS presence and asymmetry in schizophrenia, obsessive-compulsive disorder, bipolar disorder, and autism spectrum disorder (2,30–38) supporting the idea that PCGS presence, not minute differences in morphology, is more suggestive of psychiatric and neurodevelopmental vulnerability. Finally, given prior work linking sulcal depth and gyrification in other implicated regions to treatment response in ADHD (61,62), an important next step would be to test whether PCGS presence or morphology similarly predicts response to stimulant or non-stimulant pharmacotherapy, which would extend the clinical utility of this structural marker beyond diagnosis and symptom severity to treatment planning.

## Conclusion

Taken together, these findings advance understanding of how early neurodevelopmental variation in cortical folding may serve as a structural intermediary between genetic liability and the behavioral expression of ADHD. By demonstrating that reduced left PCGS presence is associated with both ADHD diagnosis and symptom severity, this study identifies a stable, early-established anatomical marker that complements and extends prior volumetric and thickness-based approaches to ADHD neuroanatomy. More broadly, these results reinforce the value of tertiary sulcal morphology as a neuroanatomical lens into individual differences in cognition and psychopathology, and they directly inform the hemispheric asymmetry hypothesis of ADHD by showing that a specific, developmentally stable structure within the ACC contributes to individual differences in ADHD symptomatology.

## Acknowledgments and Disclosures

All authors report no biomedical financial interest or potential conflicts of interest. This research was supported by NSF CAREER Award 2042251 (Weiner), Medical Scientist Training Program Grant T32 GM140935 (Willbrand), and National Institutes of Health F31AG082446 (Maboudian); funding for this project was also provided by the University of Wisconsin Department of Radiology R&D Committee (Willbrand). Funding for original data collection and curation for the New York University sample was provided by the National Institute of Mental Health (R01MH083246), Autism Speaks, The Stavros Niarchos Foundation, The Leon Levy Foundation, and An endowment provided by Phyllis Green and Randolph Cōwen. The funding agencies did not have a role in the study design, data collection and analysis, decision to publish, or preparation of the manuscript.

## Author contributions

RFH, AP, MA, EHW, and KSW designed research; RFH, AP, MA, EM, CMG, SAM, EHW, and KSW performed research; RFH, AP, MA, EM, CMG, EHW, and KSW wrote the paper; all authors gave final approval to the paper before submission.

## Data and code availability

All data and original code used for the present project will be publicly available on GitHub upon publication (https://github.com/cnl-berkeley/stable_projects). Any additional information required to reanalyze the data reported in this paper is available from the corresponding author (Kevin Weiner) upon request.

## Notes

### Competing Interest Statement

The authors have declared no competing interest.

