## Supplemental Material for "Linking anterior cingulate sulcal variability to attention-deficit hyperactivity disorder behavior"

We also sought to relate PCGS morphological features to each behavioral metric. We regressed hemispheric PCGS length, depth, and cortical thickness on ADHD index, inattentive score, and hyperimpulsivity. Of these, only the length of the LHS PCGS showed a relationship with hyperimpulsivity score, but when adjusted with False Discovery Rate (FDR) methods, this was insignificant (adjusted $R^2$ = 0.117, P = 0.007, P(FDR) = 0.122). Further linear regressions including morphological feature * diagnostic group as a feature also showed a significant uncorrected interaction between diagnosis and right hemisphere PCGS mean cortical thickness on hyperimpulsivity scores (adjusted $R^2$ = 0.564, P = 0.043). This indicates that the structural relationship between right ACC cortical thickness and impulsive behavior differs between the ADHD and neurotypical cohorts. However, this interaction did not survive multiple comparison corrections across all morphological features (P(FDR) = 0.918). No other morphological interaction terms reached statistical significance (Ps>0.05).

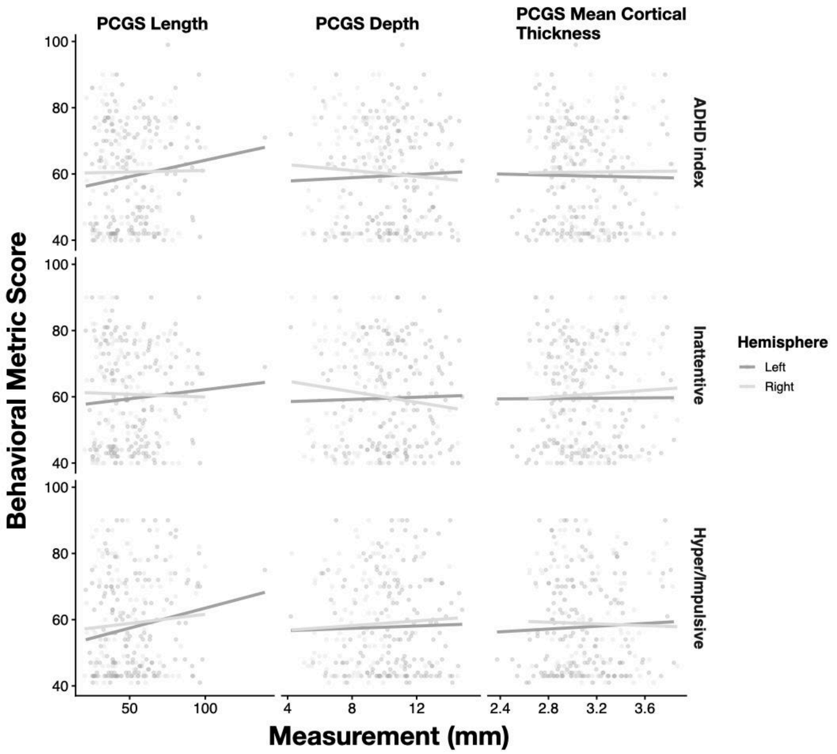


**Figure S1. Quantitative measurements of the PCGS do not correlate with ADHD-related behavioral metric scores.** Linear correlation between the length (left panel), depth (middle panel) and cortical thickness mean (right panel) and behavioral metrics (ADHD index, top; inattentiveness, middle; hyperimpulsivity, bottom). Statistical significance was derived from multivariable linear regression models controlling for age, gender, and handedness, with p-values adjusted for multiple comparisons using False Discovery Rate (FDR). The length of the PCGS in the left hemisphere was marginally associated with Hyper Impulsivity but did not survive FDR correction (p=0.0067 before correction, p(corrected)=0.122).
